# The oncogenic transcription factor FUS-CHOP can undergo nuclear liquid-liquid phase separation

**DOI:** 10.1101/2021.02.24.432743

**Authors:** Izzy Owen, Debra Yee, Hala Wyne, Theodora Myrto Perdikari, Victoria Johnson, Jeremy Smyth, Robert Kortum, Nicolas L. Fawzi, Frank Shewmaker

**Affiliations:** Department of Biochemistry and Molecular Biology, Uniformed Services University, Bethesda, MD, USA; Center for Biomedical Engineering, Brown University, Providence, RI, USA; Department of Molecular Pharmacology, Physiology, and Biotechnology, Brown University, Providence, RI, USA; Department of Anatomy, Physiology and Genetics, Uniformed Services University, Bethesda, MD, USA; Department of Pharmacology and Molecular Therapeutics, Uniformed Services University, Bethesda, MD, USA

## Abstract

Myxoid liposarcoma is caused by a chromosomal translocation resulting in a fusion protein comprised of the N-terminus of FUS (fused in sarcoma) and the full-length transcription factor CHOP (CCAAT/Enhancer Binding Protein Homologous Protein). FUS functions in RNA metabolism and CHOP is a stress-induced transcription factor. The FUS-CHOP fusion protein causes unique gene expression and oncogenic transformation. The FUS segment is required for oncogenic transformation, but the mechanism of FUS-CHOP-induced transcriptional activation is unknown. Recently, some transcription factors and super enhancers were proposed to undergo liquid-liquid phase separation and form membraneless compartments that recruit transcription machinery to gene promoters. Since phase separation of FUS depends on its N-terminus, transcriptional activation by FUS-CHOP could result from the N-terminus driving nuclear phase transitions. Here, we characterized FUS-CHOP in cells and *in vitro*, and observed novel phase-separating properties relative to unmodified CHOP. Our data indicate FUS-CHOP forms phase-separated condensates at super enhancer transcriptional sites. We provide strong evidence that the FUS-CHOP phase transition is a novel oncogenic mechanism and potential therapeutic target for treatment of myxoid liposarcoma.

## Introduction

Soft tissue sarcomas (STS) are diagnosed in roughly 12,000 patients in the United States each year with a mortality rate of approximately 40% (Siegel et al., 2015). Liposarcoma is the most common type of STS, accounting for around 20% of all adult STS diagnoses (Bock et al., 2020; Perez-Losada et al., 2000). Myxoid liposarcoma (MLS) is the second most common liposarcoma and is distinguished by the cytogenetic hallmark of t(12;16)(q13;p11) (Bock et al., 2020; Suzuki et al., 2012). This chromosomal translocation creates a novel fusion protein composed of the N-terminus of FUS (fused in sarcoma) and full-length CHOP (CCAAT/Enhancer Binding Protein (C/EBP) Homologous Protein).

FUS is a ubiquitously expressed, predominantly nuclear DNA and RNA binding protein that functions in the DNA damage response, transcription, and RNA metabolism (Chen et al., 2019a; Tan et al., 2012; Zinszner et al., 1997). The N-terminal prion-like domain of FUS (~aa1-165) (PrLD) is required for FUS self-association, chromatin binding, and transcriptional activation (Yang et al., 2014). CHOP is a member of the C/EBP family of transcription factors that play a role in differentiation, proliferation, and energy metabolism in various cell types (Hu et al., 2018). Normally, CHOP expression is suppressed, but upregulated during differentiation and following cellular stress (Ohoka et al., 2007; Yang et al., 2017). In MLS, the fusion protein, FUS-CHOP, is expressed under the control of the FUS promoter, resulting in ubiquitous expression. Importantly, ubiquitous overexpression of CHOP alone in nude mice does not result in MLS, whereas expression of FUS-CHOP from the same promoter does, indicating that FUS provides novel oncogenic properties to the fusion protein (Perez-Losada et al., 2000).

Genome wide occupancy analysis of MLS cell lines found that 60% of FUS-CHOP mapped to putative enhancers, occupying 97% of super enhancers (SEs) defined by the presence of the H3K27ac chromatin modification (Chen et al., 2019b). SEs are clusters of transcriptional enhancers that recruit a high density of transcriptional regulators and machinery (Thandapani, 2019). In cancer, SEs can contain numerous mediators, signaling factors, RNA polymerase II, and chromatin modifications (such as H3K27ac), which function together as regulators of oncogene expression (Bradner et al., 2017). In liposarcomas, SEs are involved in amplifying cancer pathways, cell migration, and angiogenesis ^12^. A recent hypothesis is liquid-liquid phase separation (LLPS) of protein activators into condensates at SEs can control gene expression (Sabari et al., 2018; Schneider et al., 2021). Such condensation of macromolecules into distinct liquid-phase states—sometimes called membrane-less organelles (MLOs) or biomolecular condensates—is attributed to many cellular functions that require spatiotemporal regulation (Banani et al., 2017; Boija et al., 2021). The results of numerous studies suggest RNA polymerase II, transcription factors, coactivators, super-enhancer sequences, mediator proteins (MED1), and other transcriptional machinery functionally undergo LLPS at transcriptional start sites (Boehning et al., 2018; Bolte and Cordelieres, 2006; Cho et al., 2018; Gurumurthy et al., 2019; Sabari et al., 2018; Wei et al., 2020).

FUS and homologous proteins, EWS and TAF15, have been shown to undergo LLPS under several conditions (Chong et al., 2018; Maharana et al., 2018; Patel et al., 2015) and recruit the C-terminal domain of RNA polymerase II into in vitro condensates (Burke et al., 2015). During transcription, FUS and RNA polymerase II are suggested to co-localize into nuclear condensates (Thompson et al., 2018). In an MLS cell line, FUS-CHOP was found to co-localize at SEs with BRD4 – a protein that is proposed to control gene expression through the formation of phase-separated condensates at SEs (Sabari et al., 2018). FUS’s N-terminal intrinsically disordered, low-complexity (LC) region (~aa 1-212) facilitates LLPS in cells and *in vitro* (Burke et al., 2015; Monahan et al., 2017), and importantly, all MLS-causing FUS-CHOP translocations contain portions of this LC sequence. Similarly, the N-terminal regions of FUS, TAF15 and EWS are all translocated in various forms of sarcomas and leukemias, including Ewing’s sarcoma, when fused to any of about a dozen transcription factors (Bolte and Cordelieres, 2006). Previous work has shown FUS-CHOP localizes to nuclear punctate structures whereas FUS and CHOP are individually diffuse nuclear proteins (Thelin-Jarnum et al., 2002). These punctate structures were eliminated by truncation of FUS’s LC region, thus restoring diffuse localization of FUS-CHOP when large segments of the LC region were removed (Goransson et al., 2002). The mechanism by which FUS-CHOP induces oncogenesis remains unknown; however, based on the above observations, we hypothesized that FUS-CHOP has novel phase-separating properties that may induce oncogenesis through condensate formation at transcription sites.

Here, we evaluate the propensity of FUS-CHOP to undergo LLPS *in vitro* and in cells. We assess localization of FUS-CHOP in MLS cancer cell lines and we demonstrate that ectopically expressed FUS-CHOP nuclear puncta have distinct liquid-like characteristics. We also observed FUS-CHOP puncta to colocalize with BRD4, which is a marker of phase-separated SEs (Sabari et al., 2018). A previous study suggested that fusion proteins with intrinsically disordered regions (IDRs) may form transcriptional condensates that induce oncogenesis (Boija et al., 2021). Likewise, our results suggest FUS-CHOP can undergo a liquid-phase transition in the nucleus, which could provide the mechanism for its emergent gain-of-function oncogenicity. This may be a general mechanism for transcriptional activation by fusion oncoproteins with IDRs.

## Results

### Recombinant FUS-CHOP undergoes LLPS *in vitro*

Both recombinant full-length FUS and its LC region have previously been shown to undergo LLPS *in vitro* (Burke et al., 2015; Patel et al., 2015). To determine if recombinant FUS-CHOP can undergo LLPS under similar conditions, we purified the most common type of FUS-CHOP translocation, type II (11 truncation variants of FUS-CHOP have been characterized from patient samples, with the most common being type II and type I, respectively (Figure S1)) (Bode-Lesniewska et al., 2007; Oikawa et al., 2012). FUS-CHOP type II contains the first 175 amino acids of FUS fused to full-length CHOP (Figure 1A). Using the N-terminal maltose binding protein (MBP) tag that we previously used for full-length FUS (Burke et al., 2015; Monahan et al., 2017; Owen et al., 2020), we purified FUS-CHOP type II and CHOP (as a control) from *Escherichia coli* (attempts to purify FUS-CHOP type I were not successful due to insolubility). Similar to our previous observations for wild-type full-length FUS (Burke et al., 2015; Monahan et al., 2017; Owen et al., 2020), upon cleavage of the MBP tag with TEV protease, we observed FUS-CHOP type II droplet formation (Figure 1B). Concomitant with phase separation, we observed increased turbidity over time for FUS-CHOP type II (Figure 1C). CHOP alone showed no droplets or marked turbidity after cleavage of the MBP tag, demonstrating that elements within FUS’s LC sequence drive phase separation. These data suggest that the LC domain of FUS provides FUS-CHOP a greater capacity to self-associate and undergo LLPS relative to the unfused CHOP protein.

**Figure 1.**
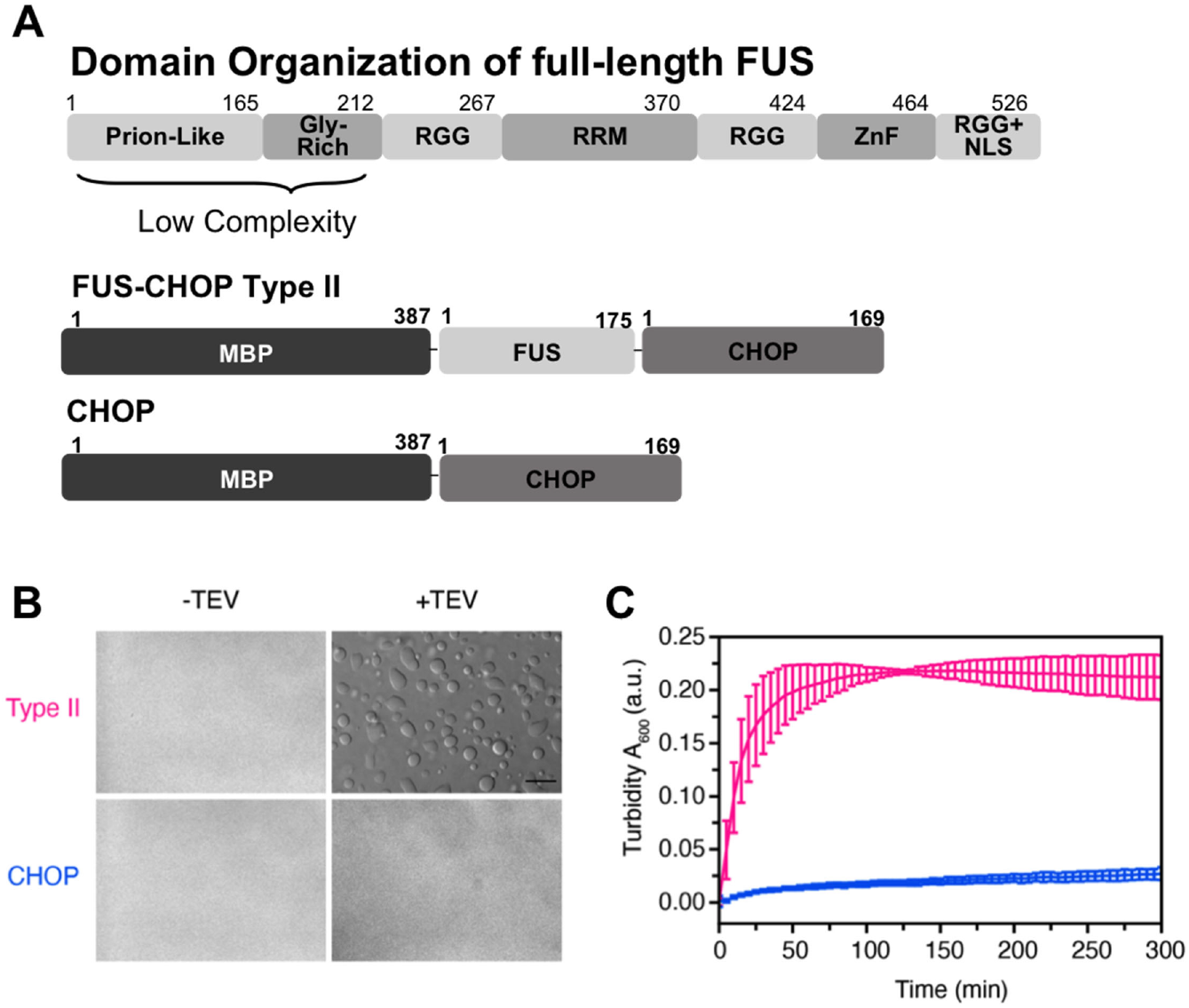
FUS-CHOP Type II undergoes LLPS *in vitro*. A–Schematic of full length FUS as well as purified recombinant MBP-FUS-CHOP and MBP-CHOP. B–DIC micrographs of 50 μM FUS-CHOP Type II fusion (top) and CHOP protein alone (bottom) in 50 mM Tris, 150 mM NaCl, pH 7.4 without and with TEV protease to cleave the N-terminal maltose binding protein (MBP) solubilizing tag. Scale bar represents 80 μm. C–Corresponding turbidity measurements of FUS-CHOP type II fusion (pink) and CHOP protein alone (blue) after initiating cleavage of the MBP tag by addition of TEV protease. Error bars represent the standard deviation of measurements from three replicates.

### Ectopically expressed FUS-CHOP-eGFP is undergoing LLPS in the nucleus

Previous work showed ectopically expressed FUS-CHOP-GFP type II formed distinct nuclear puncta (Thelin-Jarnum et al., 2002). To confirm this observation, we ectopically expressed FUS-CHOP type I and type II eGFP-tagged fusion proteins, in NIH 3T3 cells (Figure 2A,B). We observed numerous round nuclear puncta of both the type I and type II constructs (Figure 2C). To ensure these structures were not the result of the eGFP tag, we also expressed untagged FUS-CHOP type I and type II (figure 2B); the untagged proteins formed similar punctate structures (Figure 2C). This nuclear punctate localization pattern of FUS-CHOP is not diffuse like either wild-type FUS or CHOP (Thelin-Jarnum et al., 2002); thus we hypothesized the puncta are phase-separated condensates driven by FUS’s intrinsically disordered LC region.

**Figure 2.**
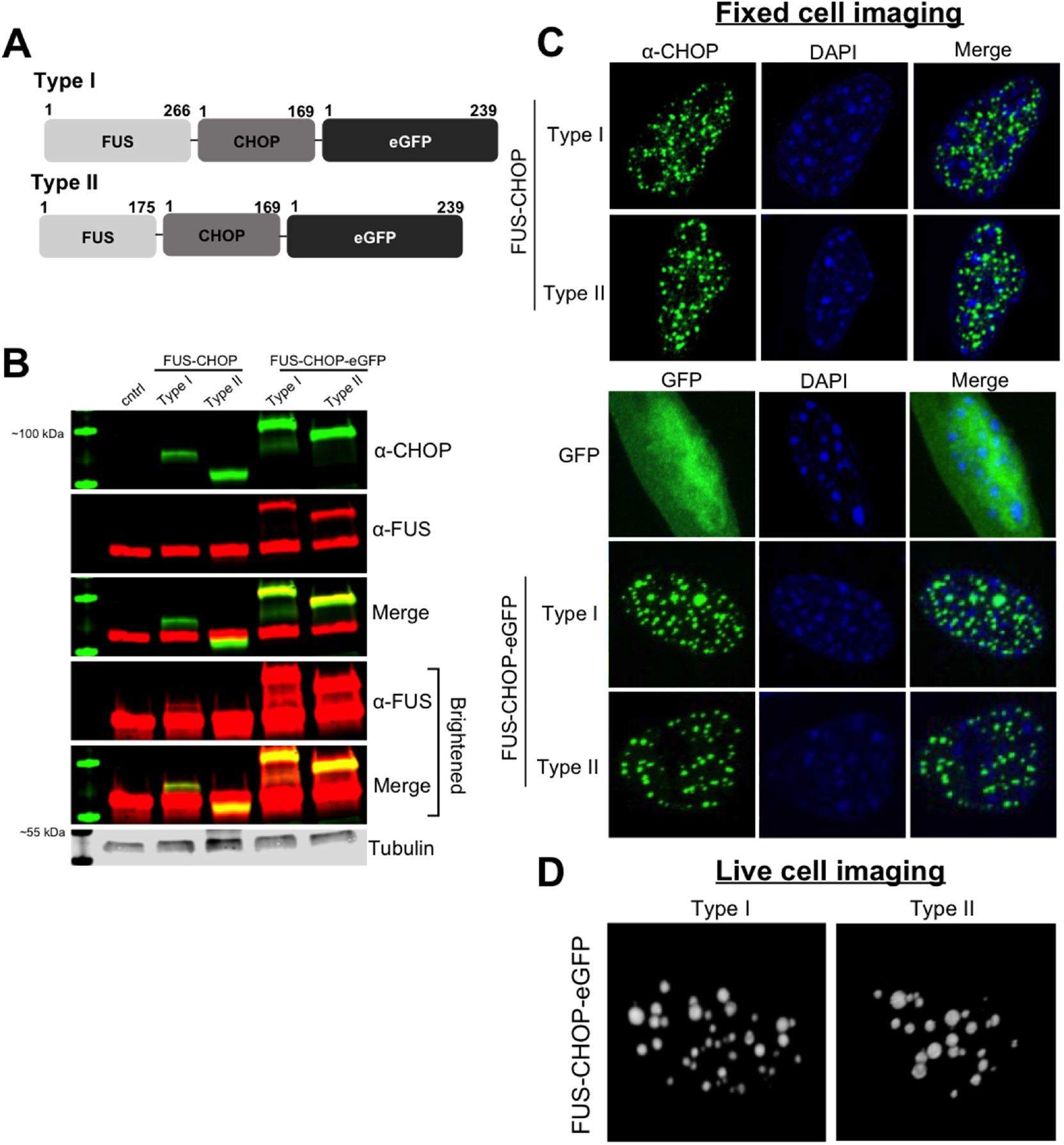
Ectopically expressed FUS-CHOP localizes to sphere shaped puncta in the nucleus. A– Schematic of FUS-CHOP-eGFP fusion proteins. B–Western Blot of ectopically expressed untagged FUS-CHOP and FUS-CHOP-eGFP in NIH 3T3 cells. The bottom panels have been brightened to show all FUS antibody binding. C–Confocal images of nuclear puncta formed by ectopically expressed FUS-CHOP (with and without eGFP tag) in NIH 3T3 cells. D–FUS-CHOP-eGFP puncta are spherical in shape when assessed in 3-Dimension. Representative data from three experimental replicates.

Phase-separated condensates in cells are typically characterized by three hallmarks: spherical in shape, fusing upon touching, and rapid internal dynamics and external exchange (Hyman et al., 2014). We used live-cell imaging to assess the puncta in 3-Dimensions. We observed spherical nuclear puncta of FUS-CHOP-eGFP type I and type II (Figure 2D). Imaging the cells over time revealed free movement around the nucleus consistent with Brownian motion. We observed frequent fusion events in which a single sphere formed when two spheres made contact (Figure 3, Figure S2, Mov 1). To quantify internal rearrangement and external exchange of FUS-CHOP puncta, we used fluorescence recovery after photobleaching (FRAP). Previous work indicates that intracellular liquid-state condensates have half-times of recovery from seconds to minutes(Banani et al., 2017). Here, we bleached both type I and II puncta and observed an average half-time of recovery of ~19 seconds and ~14 seconds, respectively (Figure 4). These data show that ectopic FUS-CHOP forms nuclear condensates and has the major hallmarks of LLPS.

**Figure 3.**
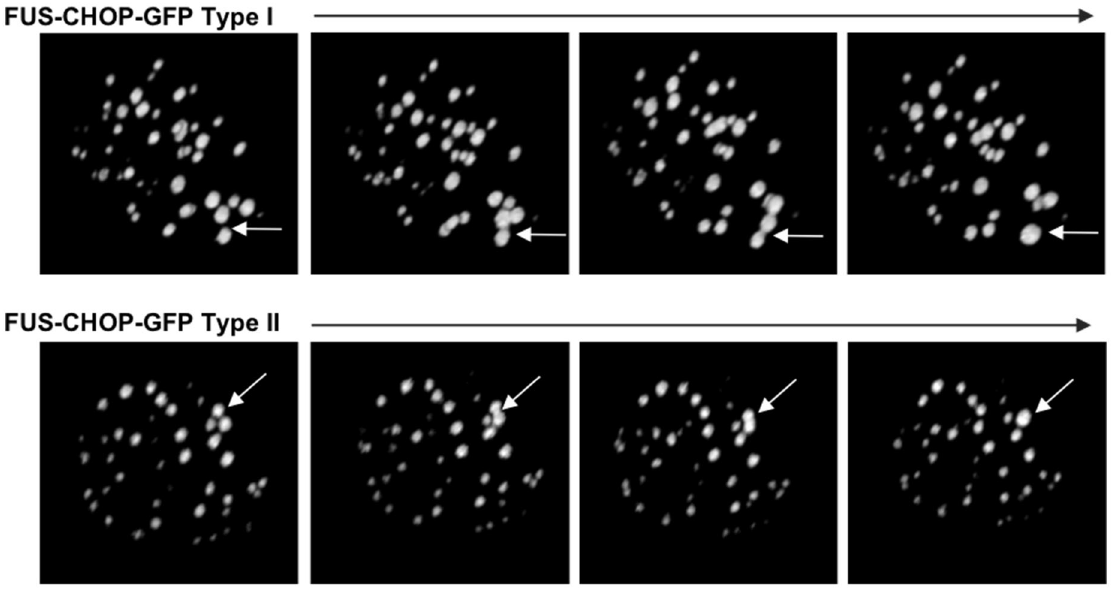
FUS-CHOP-eGFP puncta have liquid-like characteristics and undergo fusion in the nucleus. Still frames from time course movies imaged by confocal microscopy of FUS-CHOP-eGFP type I and type II puncta fusing upon touching. Representative data from three experimental replicates. Videos are available in Supplementary Data.

**Figure 4.**
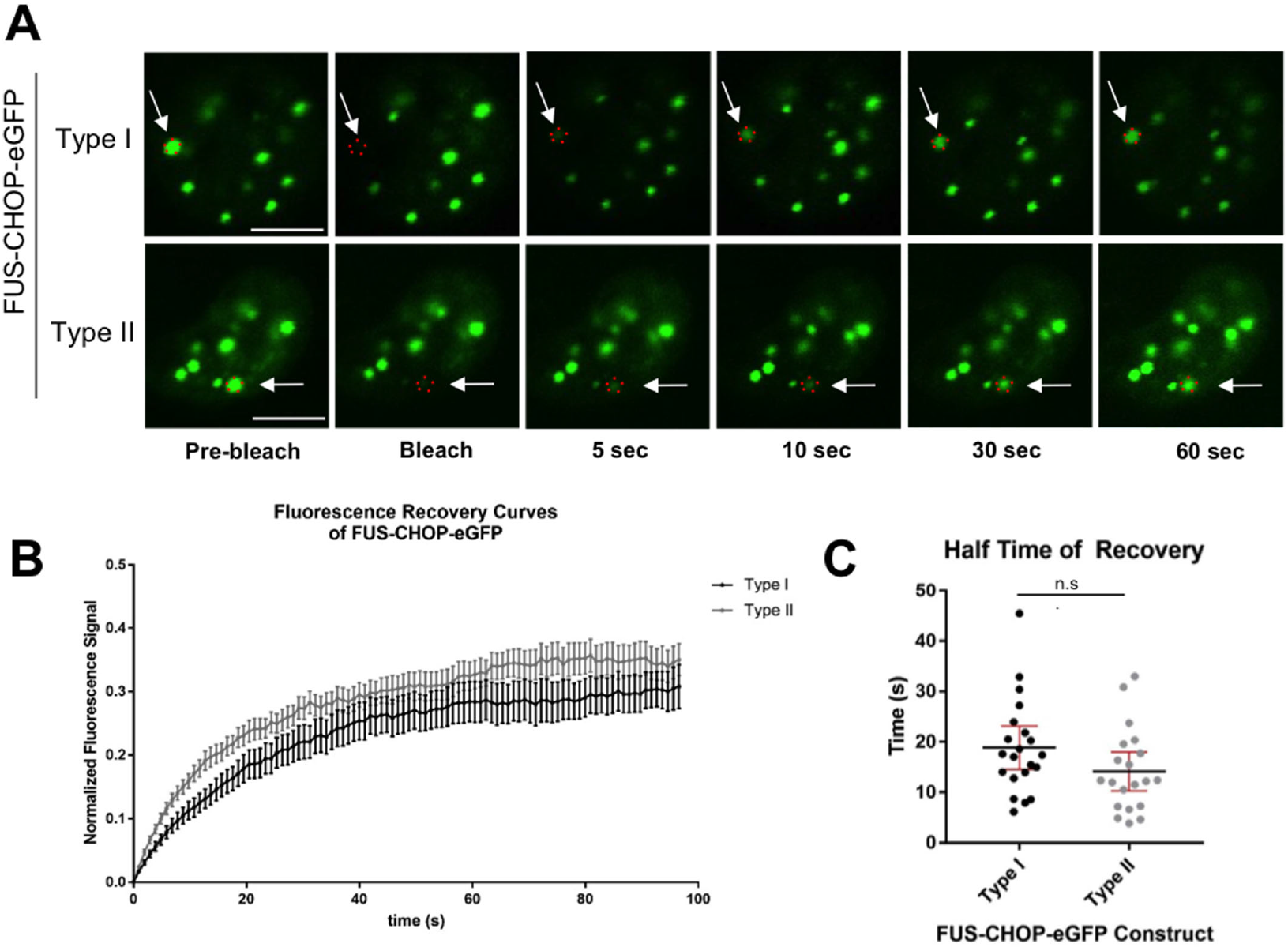
FUS-CHOP-eGFP has liquid-like recovery following fluorescence bleaching. A–FUS-CHOP-eGFP type I and type II puncta recover on the time scale of seconds following fluorescence bleaching. Scale bar represents 5 μm. Representative data from three experimental replicates. B– Average fluorescence recovery curves of FUS-CHOP-eGFP type I and type II. C–Half time of recovery of FUS-CHOP-eGFP type I and II puncta. Error bars represent the mean with 95% c.i. of measurements from three replicates (B,C).

We also observed that the type I puncta moved more rapidly (they were less static) than the type II. We hypothesize that the RGG repeats present in the longer type I fusion (but not the type II; Figure 1A, 2A) could be driving additional protein-protein (Ryan et al., 2018; Wang et al., 2018) or protein-RNA interactions (as the LC does not interact with RNA (Burke et al., 2015)), leading to more mobility throughout the nucleus. However, we did not pursue this issue further.

### Phase separation of FUS-CHOP-eGFP is dependent on the FUS prion-like domain

FUS’s LC domain is composed of a PrLD domain followed by a short glycine-rich region (Figure 1A) – both of which are IDRs with little complexity in their amino-acid composition. PrLDs are frequently linked to both LLPS and formation of pathological inclusions in neurodegenerative diseases (March et al., 2016). FUS’s PrLD facilitates LLPS of wild-type FUS (Burke et al., 2015), but its truncation inhibits LLPS (Patel et al., 2015). In earlier work when the PrLD of FUS-CHOP fusions was serially truncated, there was a concomitant dissolution of nuclear puncta (Goransson et al., 2002). To determine the dependence of phase separation on the PrLD (~aa 1-165), we removed the first 25, 50, 75 and 125 amino acids in type I and type II fusion constructs of FUS-CHOP-eGFP and expressed them in NIH 3T3 cells (Figure 5A, B). We observed retention of punctate structures upon removal of up to 50 amino acids (type 1 is shown in Figure 5C; identical results for type II are in Figure S3). Removing 75 or 125 amino acids resulted in a diffuse pattern of expression (Figure 5C, Figure S3).

**Figure 5.**
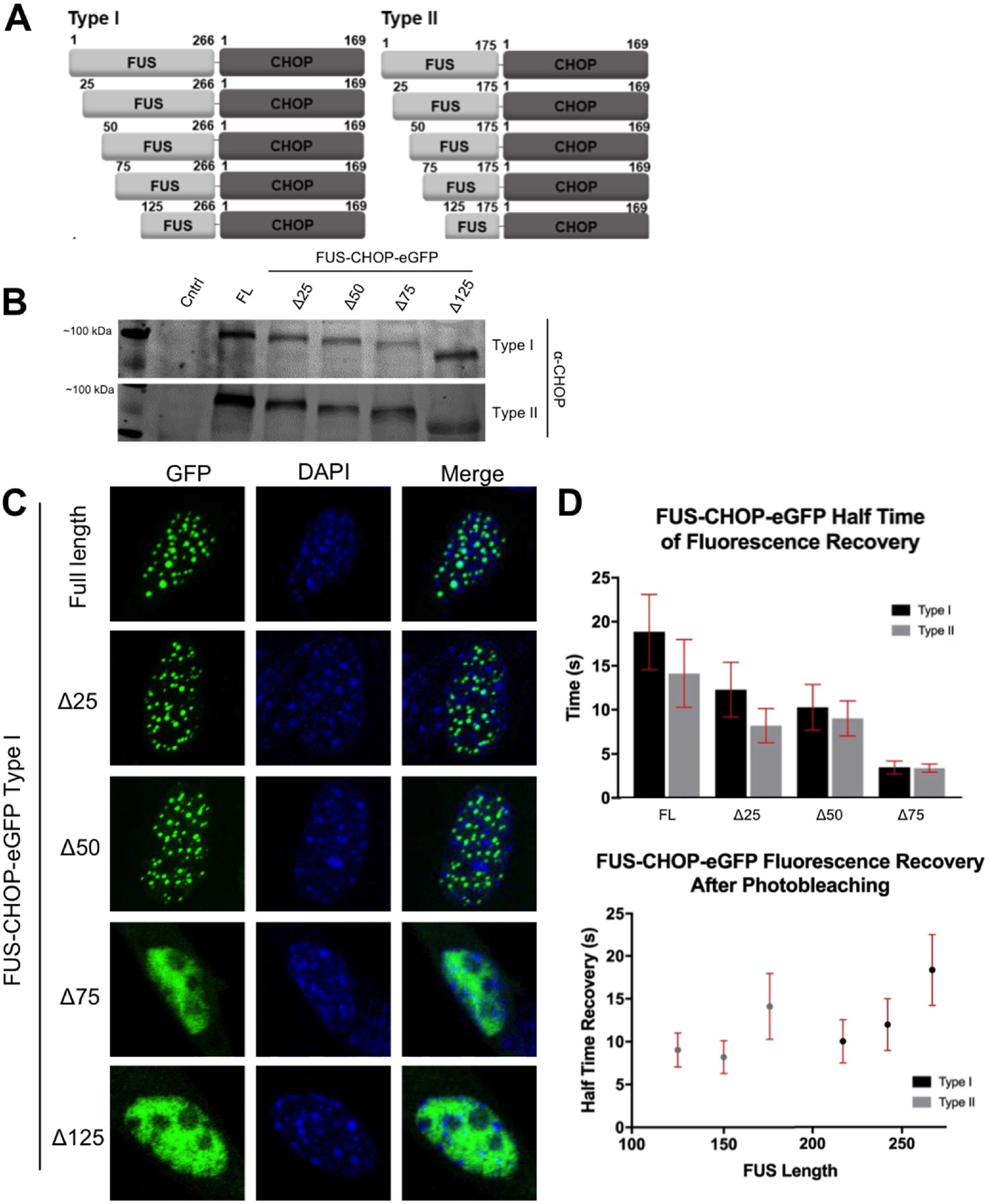
FUS-CHOP-eGFP liquid-liquid phase separation is dependent on the N-terminus of FUS. A–Schematic of truncations made to FUS-CHOP-eGFP type I and type II. B–Western blot of NIH 3T3 cells transfected with full-length or truncated FUS-CHOP-eGFP constructs. NIH 3T3 cell lysates were probed with anti-CHOP antibody. C–Full-length or truncated FUS-CHOP-eGFP type I ectopically expressed in NIH 3T3 cells and imaged by confocal microscopy. Representative data from three experimental replicates. Type II images are shown in Supplementary Data. D–FUS-CHOP-eGFP half-life of recovery is dependent on the length of FUS’s prion-like domain. Error bars represent the mean with 95% c.i. of measurements from three replicates.

The above observation suggests that the interactions driving FUS-CHOP phase separation require most of the PrLD to be intact (See Figure 1A). These data could point to a special feature or structure in the region spanning residues 51 to 75 of the FUS PrLD. Yet, our previous work suggests that the PrLD does not populate specific structures even in the liquid form (Murthy et al., 2019). Therefore, we also created internal PrLD truncations to test the hypothesis that the location of the truncation is not important, but instead the total length of the PrLD present determines phase separation. To this end, we deleted amino acids 50-75 in both FUS-CHOP-eGFP type I and type II constructs (Figure 6A) and ectopically expressed them in NIH 3T3 cells to determine their effects on LLPS (Figure 6B,C). As with the N-terminal truncations, these constructs were observed to form spherical nuclear punctate structures (Figure 6B,C). Therefore, these data suggest that the length and not the location of the PrLD deletion determines in-cell phase separation of FUS-CHOP-eGFP.

**Figure 6.**
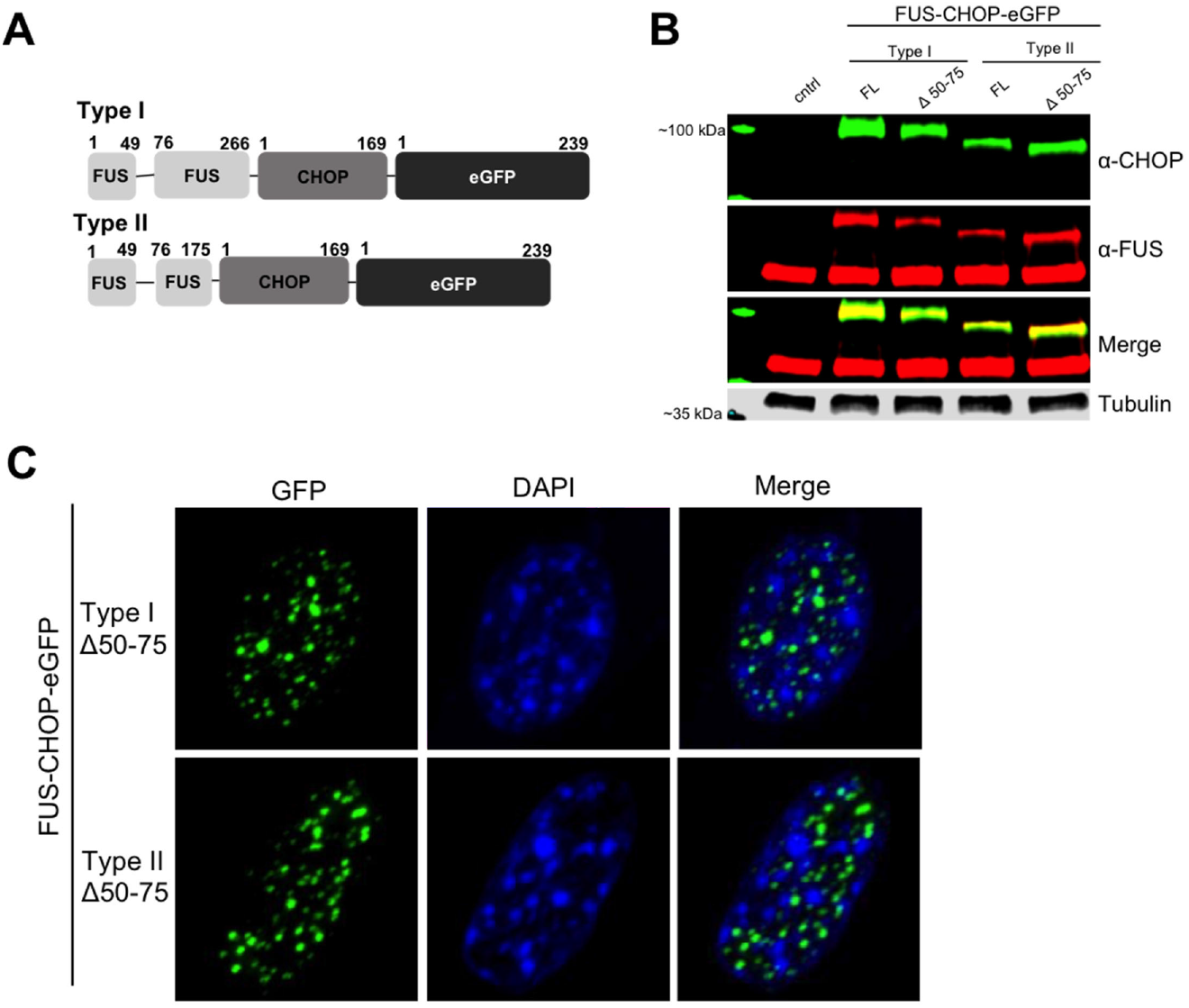
Phase separation of FUS-CHOP is not dependent on a central core region within FUS’s prion-like domain. A–Schematic of FUS-CHOP-eGFP type I and type II internal truncations. B–NIH 3T3 cells transfected with full-length (FL) or truncated (Δ 50-75) FUS-CHOP-eGFP type I or type II constructs. Cell lysates were analyzed by Western Blot and probed with anti-CHOP and anti-FUS antibodies. C– Confocal images of internally truncated FUS-CHOP-eGFP nuclear puncta type I or type II. Representative data from three experimental replicates.

To characterize how the PrLD of FUS affects FUS-CHOP LLPS, we used FRAP to quantify recovery time of the truncated proteins within the puncta. We observed a quicker half time of recovery as the PrLD was shortened, suggesting length can affect dynamic movement into or within the phase-separated condensate (Figure 5D). Type I condensates contain more of FUS’s N-terminal sequence (266aa) and consistently recover slower than type II (175aa), including truncated proteins. The length and low-complexity features of FUS’s PrLD appear to be the dominant factors in governing FUS-CHOP LLPS, as opposed to any particular sequence element.

### FUS-CHOP is localized in small nuclear punctate structures in myxoid liposarcoma cell lines

We next sought to characterize endogenous FUS-CHOP in patient-derived cells. We assessed endogenous expression and localization of FUS-CHOP in three different MLS cell lines. MLS-402 and MLS-1765 were both established and immortalized by transfection with SV40 large T-antigen, while DL-221 was spontaneously immortalized from patient tumor samples (Aman et al., 1992; de Graaff et al., 2016; Thelin-Jarnum et al., 1999). MLS-402 and DL-221 both contain type I fusions like we used in ectopic expression, while MLS-1765 has a type VIII fusion that encompass the first 514aa of FUS (Figure 7A; Figure S1). All three cell lines showed FUS-CHOP localized to small nuclear punctate structures, similar to those seen in a previous study evaluating oncogenic EWS-FLI1 fusions in cancer cell lines (Chong et al., 2018) (Figure 7B, Mov 2). Localization of FUS-CHOP in all cell lines was punctate and nuclear, but expression levels of type VIII were greater than type I (Figure 7C). Fixed-cell imaging indicated smaller punctate structures in the cancer cell lines than observed for the ectopically expressed proteins in NIH 3T3 cells.

**Figure 7.**
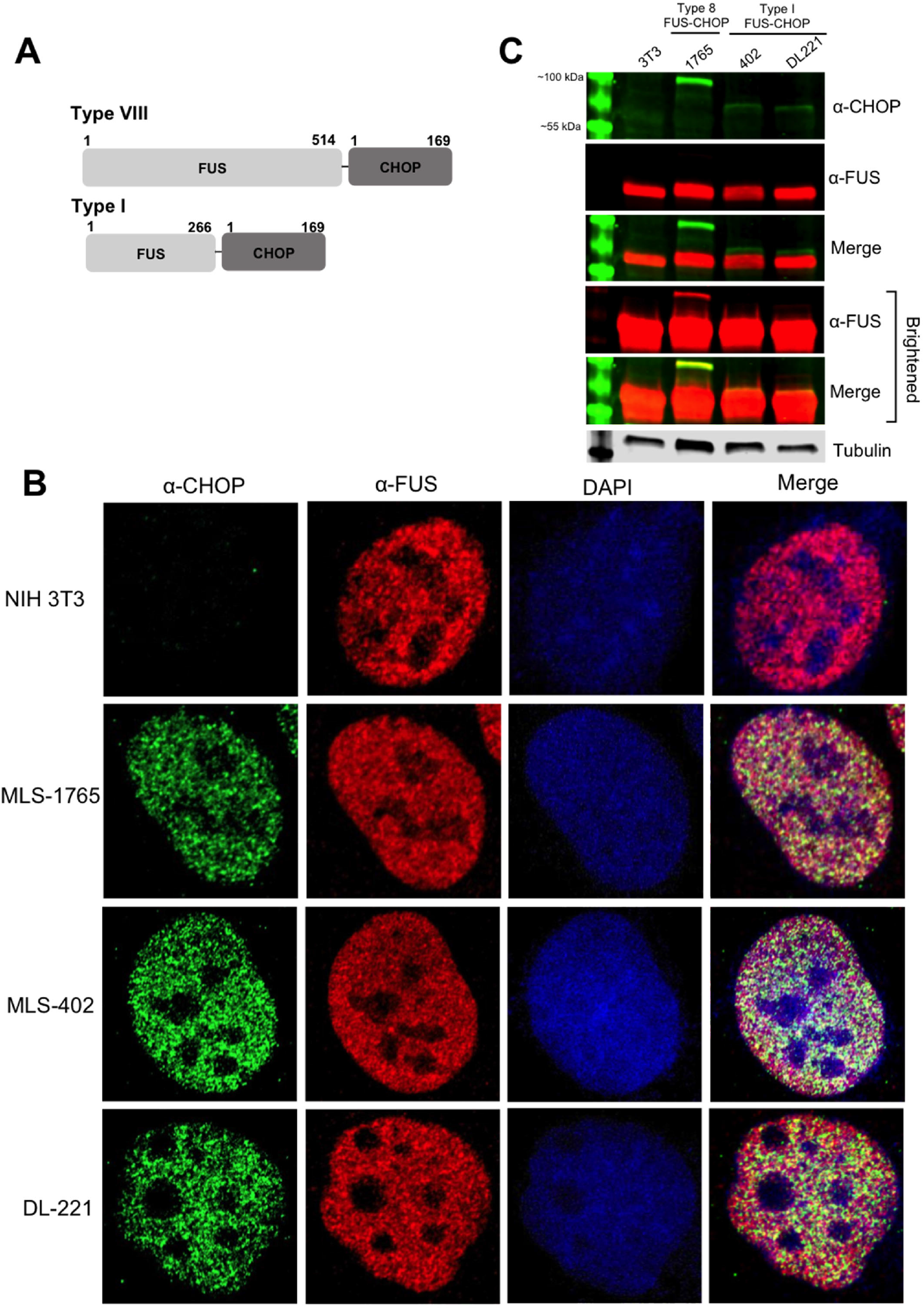
FUS-CHOP forms nuclear puncta in Myxoid Liposarcoma cell lines. A–Schematic of FUS-CHOP fusions. MLS-1765 encodes a type VIII FUS-CHOP fusion protein while MLS-402 and DL-221 contain a type I fusion protein (diagram in Supplementary Figures). B–MLS cell lines were probed with anti-CHOP and anti-FUS antibodies and imaged using confocal microscopy with Airyscan. C–Cancer cell lysates analyzed by Western Blot. Brightened Blot shows FUS antibody binding. Representative data from three experimental replicates.

### FUS-CHOP is localized to BRD4 phase separated puncta

In a previous study, FUS-CHOP was shown to occupy 9% of active promoter sites and 60% of putative enhancer sites in an MLS cell line (Chen et al., 2019b). Using ChIP-seq, the authors found that FUS-CHOP occupied 40% of the same enhancers as BRD4 (Chen et al., 2019b), which itself localizes to enhancer sites marked by acetylated histones (Loven et al., 2013). The authors concluded that FUS-CHOP and BRD4 cooperate at oncogenic SEs in MLS (Chen et al., 2019b). Importantly, BRD4 has an intrinsically disordered C-terminal domain that purportedly drives its phase separation into nuclear puncta at super enhancers in cell models (Sabari et al., 2018). If FUS-CHOP is undergoing LLPS at SEs in our MLS cell lines, then we would predict colocalization with BRD4 at nuclear puncta.

We probed our MLS cancer cell lines for BRD4 puncta and assessed its colocalization with FUS-CHOP (Figure 8A). Pearson’s correlation coefficient between BRD4 and FUS-CHOP was 0.705, 0.727, and 0.691 in MLS-1765, MLS-402, and DL221 cell lines, respectively. We also evaluated colocalization in our ectopic-expression model. We expressed both FUS-CHOP-eGFP type I and type II in NIH 3T3 cells and probed for BRD4 (Figure 8B). We saw small BRD4 puncta throughout the nucleus in the control, but in the FUS-CHOP-GFP expressing cells, BRD4 localized to the large FUS-CHOP-eGFP puncta. These data suggest FUS-CHOP and BRD4 occupy the same nuclear condensates at SEs.

**Figure 8.**
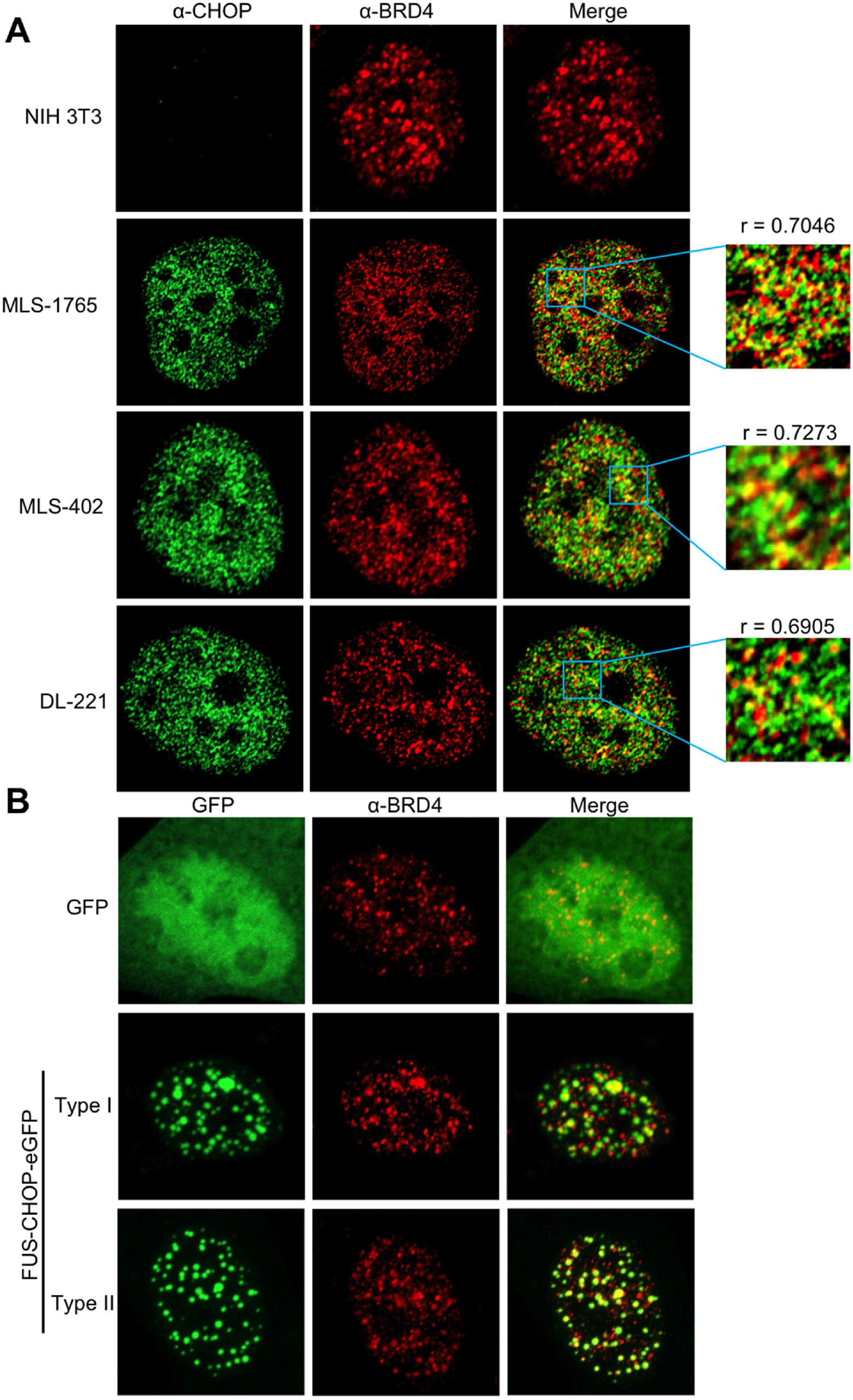
FUS-CHOP localizes with phase-separating super-enhancer protein BRD4. A–MLS-1765, MLS-402, and DL-221 cells were probed with anti-CHOP and anti-BRD4 antibodies and analyzed by confocal microscopy with Airyscan. Pearson’s correlation coefficient was calculated using the JaCoP plug-in in Fiji ^56^. B–NIH 3T3 cells were transfected with FUS-CHOP-eGFP type I or type II for 24 hours. Following transfection, cells were probed with anti-BRD4 and assessed by confocal microscopy with Airyscan. Representative data from three experimental replicates.

## Discussion

Macromolecular condensates that form via LLPS are implicated in many sub-cellular processes (Alberti and Hyman, 2021). Condensates have recently been proposed to also have roles in pathological events like oncogenic transcription (Boija et al., 2021). In such cases, oncogenesis would depend on the condensation of transcription factors, mediator complex proteins, and chromatin remodeling proteins to high density at specific SEs and promoter sequences (Boija et al., 2021). Here, we assessed the pathological FUS-CHOP fusion protein—which causes aberrant transcription in MLS (Joseph et al., 2014) — and its capacity to undergo LLPS *in vitro* and in cell models. Our data indicate that N-terminal regions of FUS provide CHOP with novel LLPS properties. In MLS cell lines, we observed the localization of FUS-CHOP condensates at SEs, suggesting that FUS-CHOP LLPS at transcriptional start sites could be integral to oncogenic mechanisms.

FUS is a member of the FET family of proteins, along with EWS and TAF15, which can all undergo LLPS in the nucleus of cells and *in vitro* (Maharana et al., 2018). All three proteins contain an intrinsically disordered, N-terminal PrLD responsible for driving LLPS and each protein has been found in oncogenic fusions with transcription factors (Linden et al., 2019; Riggi et al., 2007). Under endogenous conditions, the FET family proteins and their fusion partners are mostly diffuse in the nucleus (Andersson et al., 2008). However, as oncogenic fusion proteins, all localize to distinct nuclear puncta (Chong et al., 2018; Thelin-Jarnum et al., 2002). The formation of dysregulated condensates, especially at SEs, is proposed to be an underlying feature of some cancers (Boija et al., 2021). We show FUS-CHOP can undergo LLPS in the nucleus and form distinct punctate structures with liquid-like dynamics. These condensates could provide an enhanced transcriptional advantage to MLS cells. This mechanism could be common to all FET-fusion oncogenic proteins.

The fusion of EWS and the transcription factor FLI1 causes Ewing’s Sarcoma (Chong et al., 2018). The EWS-FLI1 fusion protein forms condensate-like hubs that are necessary for driving oncogenic transcription (Chong et al., 2018). Here, we see similar results with FUS-CHOP phase separation in the MLS cancer cell lines. To understand how FUS-CHOP might modify the transcriptional landscape in MLS, we looked at the localization of BRD4 – a protein shown to phase separate at SEs. BRD4 has also been shown to colocalize with FUS-CHOP in MLS (Chen et al., 2019b). The C-terminal domain that drives

BRD4’s LLPS is necessary for its function (Wang et al., 2019), suggesting that function could be linked to LLPS. Here, in MLS cancer lines, we observed FUS-CHOP nuclear condensates to localize with BRD4. Similarly, in the ectopic expression system, we observed BRD4 localization and consolidation into the large condensates composed of FUS-CHOP-eGFP. This suggests FUS-CHOP could be driving phase separation of BRD4 at oncogenic SEs in MLS. These findings provide a mechanism by which oncogenic fusion proteins, such as EWS-FLI1 and FUS-CHOP, hijack BRD4 and other bromodomain-containing proteins to induce oncogenic SEs (Chen et al., 2019b; Gollavilli et al., 2018). FUS-CHOP has also been reported to localize to sites of chromatin remodeling, specifically interacting with the SWI/SNF chromatin remodeling complex. This interaction is dependent on FUS’s PrLD, as truncation eliminates the association (Yu et al., 2019). Together, these data suggest a role of FUS-CHOP LLPS in chromatin remodeling and transcription, conferring an oncogenic advantage.

All FUS-CHOP fusion variants that cause MLS contain segments of FUS’s PrLD (many also contain longer segments that include the entire low-complexity (LC) region, but no fusions lack PrLD segments (Figure S1). The PrLD drives LLPS of full-length FUS *in vitro* and in cells (Burke et al., 2015; Wang et al., 2018). Our data indicate that the PrLD confers its phase-separating capacity to FUS-CHOP. The PrLD is approximately 165-residues long, but the induction of LLPS of FUS-CHOP type I and II could still be achieved after truncating the first 50 amino acids from the N-terminus. Since the PrLD consists mostly of a few redundant amino acids (SQGY), it does not appear that any sequence feature or motif is required to induce LLPS, but simply a segment of sufficient length. Nearly all characterized FUS-CHOP variants (10 of 11) contain the entire PrLD (Figure S1) (Oikawa et al., 2012), suggesting that shorter fusions are either less probable or less likely to induce MLS transformation.

The atomic-level structure of condensates appears to be non-static (Burke et al., 2015). However, rigid amyloid-like interactions have been proposed to support the architecture of condensates (Kato et al., 2012; Murray et al., 2017). A segment within FUS’s PrLD (residues 39-95), forms a highly ordered amyloid structure in the recombinant protein (Murray et al., 2017). It was suggested that similar amyloid-like interactions formed by this segment could underlie the structure of the phase-separated state. However, when we deleted an internal portion of the PrLD (residues 75-100), ectopic FUS-CHOP still displayed LLPS properties. This observation suggests that LLPS is a feature that emerges from the low-complexity, intrinsically disordered nature of the PrLD and is not the consequence of a precise sequence element.

Phase separation as a mechanism of transcriptional regulation has not been definitively established (McSwiggen et al., 2019); largely because transcriptional activation sites are small and dynamic, and thus make it challenging to design experiments that strongly support or refute an LLPS hypothesis (McSwiggen et al., 2019). Regardless, LLPS of enhancer-binding proteins, transcription factors and RNA polymerase II at transcriptional sites has been proposed by several groups (Boija et al., 2018; Cho et al., 2018; Hnisz et al., 2017). An LLPS model is attractive because it could explain the low-complexity, intrinsically disordered sequences that are common to transcription factors and coactivators, since these sequences can support multivalent interactions and are prone to phase-separation. Our data do not provide an answer to the molecular-level details of FUS-CHOP-induced transcription in cancer cells. However, our data clearly show that both recombinant and ectopically expressed FUS-CHOP have the capacity to undergo LLPS, whereas this property is not observed for wild-type CHOP under identical conditions. Ectopic expression is imperfect because it may cause proteins to exceed critical concentrations that would not be normally achieved in vivo (McSwiggen et al., 2019). If proteins like BRD4 are indeed marking distinct liquid-phase states at transcriptional start sites, then FUS-CHOP’s colocalization and capacity to undergo LLPS suggests that oncogenic transcription patterns could emerge from a phase-separated state. Recently, some cancer drugs have been shown to partition into biomolecular condensates (Klein et al., 2020), and drug concentration within condensates has been shown to influence therapeutic efficacy (Klein et al., 2020). If FUS-CHOP LLPS is integral to oncogenic cellular reprogramming, then this provides a new avenue for pharmacological exploration.

## Methods

### Cell culture

NIH 3T3 (ATCC CRL-1658) were cultured in DMEM (Sigma D6429) supplemented with 10% calf bovine serum (ATCC 30-2030) and 1% penicillin-streptomycin (ThermoFisher 15140148). DL-221 (MD Anderson cell core) cells were cultured in DMEM (ThermoFisher 11875093) supplemented with 10% fetal bovine serum (Sigma F6178) and 1% penicillin-streptomycin. MLS402-91 and MLS1765-92 *(received from Pierre Aman)* were cultured in RPMI supplemented with 10% fetal bovine serum and 1% penicillin-streptomycin. Cells were lysed with a modified RIPA buffer (200 mM NaCl, 100 mM Tris-HCl pH 8, 0.5% sodium deoxycholate, 1% Triton X-100, 670 mM phenylmethylsulfonyl fluoride, 1250 units of benzonase nuclease (Sigma E8263), 150 μL protease inhibitor cocktail (Thermo 1861278), and 100 μL phosphatase inhibitor (Thermo 78426)) for 30 minutes on ice.

### Transfections

DNA was transfected into NIH 3T3 cells at ~70-80% confluency using Lipofectamine 2000 (Thermo 11668027) and OptiMEM (Gibco 31985070) in a ratio of 3-6 μg DNA to 2.5 μL Lipofectamine 2000 and incubated at 37 C for 24 hours unless otherwise stated.

### Cloning/plasmids

FUS-CHOP type I and type II genes were synthesized by Genscript (Piscataway, NJ) and subcloned into pcDNA3-EGFP (Addgene 13031) or 6xHis-MBP-FUS (Addgene 98651) to produce the fusion plasmids. The FUS-CHOP truncations (Δ25, Δ50, Δ75, Δ125, and internal FUS 50-75 deletion) were generated through PCR cloning using either BamHI/Xho or HindIII/BamHI restriction sites (Thermo F531S).

Primer sequences used in the truncations were as followed:

FUS forward (CACAAGCTTATGGCCTCAAACGATTATACCCAA),

FUS Δ25 forward (CACGGATCCATGTATTCCCAGCAGAGCAG),

FUS Δ50 forward (CACGGATCCATGTATGGCCAGAGCAGC),

FUS Δ75 forward (CACGGATCCATGTATGGCTCGACTGGC),

FUS Δ125 forward (CACGGATCCATGCCCCAGAGTGGGAGC),

FUS Δ50 reverse (GTGGGATCCGCCTGAAGTGTCCGTGGA),

eGFP reverse (TGCTCACCATCTCGAG)

### Western blotting

Lysates were mixed with 4x NuPAGE LDS Sample Buffer (ThermoFisher NP0008) and electrophoresed through AnyKD precast gels (BioRad 4569034) at 80 V for 2 hours. Gels were transferred through eBlot L1 (GenScript L00686) onto nitrocellulose membranes (BioRad 1620112). Membranes were blocked with 6% milk (BioRad 1706404) in Tris buffered saline (TBS) (VWR J640). Primary and secondary antibodies were diluted in TBS with 0.1% Tween-20 (Sigma P7949). The following primary antibodies were used to probe the blots: FUS (Bethyl A300-302A), CHOP (Cell Signaling 2895S), gamma tubulin (Sigma T6557). Primary antibodies were detected with secondary antibodies conjugated to IRDye fluorescent probes (LI-COR 926-68021,926-32210). Blots were imaged with the Odyssey CLx Imaging System (LI-COR). Blot processing was done using Image Studio software (Li-COR).

### Microscopy (fixed & live cell imaging techniques)

For fixed cell imaging, cells were grown on glass coverslips for 24 hours and fixed with 4% paraformaldehyde (Sigma P6148). The cells were permeabilized in cold methanol (−20 C) and blocked with 5% normal goat serum (Abcam ab7481) with 0.05% sodium azide (Life Technologies 50062Z). The following antibodies were used to probe the fixed cells: FUS (Bethyl A300-302A), CHOP (Cell Signaling 2895S), and BRD4 (Abcam ab128874). Secondary antibodies used to detect primary antibodies were AlexaFluors AF488 and AF568 (ThermoFisher A-11001, A-11011). Nuclei were stained using Prolong mounting media with DAPI (ThermoFisher P36931). Slides were imaged using the Nikon A1R and the Zeiss 980 with Airyscan. Airyscan images were taken using the smart setup settings. Images were directly processed using the Zeiss system. All fixed cell images were further processed using ImageJ and Photoshop. Pearson’s correlation coefficient was calculated using the JaCoP plug-in in Fiji (Bolte and Cordelieres, 2006).

For live cell imaging, cells were grown in glass bottom microwell dishes 24 hours prior to transfection. The cells were incubated at 37 C for 24 hours post transfection. Before imaging, the medium was changed to dye-free DMEM (Thermo #21063029). To analyze dynamics and fusion events of FUS-CHOP spheres, time-lapse, 3-dimensional confocal imaging was carried out using the resonant scanner and Piezo Z-stage controller of the Nikon A1R microscope. Z-stacks with an interval of 0.5 μm that encompassed the nucleus of a single cell were acquired every 2 seconds over a 4 minute time period. The Z-stacks were then processed to generate 3-dimensional renderings using Nikon Elements software, and time lapse renderings were converted to video files. FRAP experiments were also carried out on the Nikon A1R. The center of a granule, marked by a 0.3 μm region of interest, was bleached at 50% power for 1.9 seconds using the 488 nm laser. The recovery was analyzed for 98 seconds (~1.5 minutes) with image acquisitions every second. The recovery was quantified using the time series analyzer V3 plugin on Fiji. The bleached pixel intensity was subtracted from each data point and then data points were normalized to the pixel intensity before the bleaching occurred. FRAP half-time data was statistically analyzed using an upaired t-test (Figure 4C, P value = 0.0953).

### *In vitro* expression and purification of FUS-CHOP fusion and CHOP

N-terminally MBP-tagged (pTHMT) FUS CHOP fusion type II and CHOP oncogene were expressed in *Escherichia coli* BL21 Star (DE3) cells (Life Technologies). Bacterial cultures were grown to an optical density of 0.7–0.9 before induction with 1 mM isopropyl-b-D-1-thiogalactopyranoside (IPTG) for 4 h at 37 C. Cell pellets were harvested by centrifugation and stored at −80 C. Cell pellets were resuspended in approximately 20 mL of 20 mM sodium phosphate, 1M NaCl, 10 mM imidazole, pH 7.4 with one EDTA-free protease inhibitor tablet (Roche) for approximately 2 g cell pellet and lysed using an Emulsiflex C3 (Avestin). The lysate was cleared by centrifugation at 47,850 *g* for 50 min at 4 C, filtered using a 0.2 μm syringe filter, and loaded onto a HisTrap HP 5 mL column. The protein was eluted with a gradient from 10 to 300 mM imidazole in 20 mM sodium phosphate, 1 M NaCl, pH 7.4. Fractions containing MBP-tagged FUS-CHOP fusion type II or CHOP were loaded onto a HiLoad 26/600 Superdex 200 pg column equilibrated in 20 mM sodium phosphate, 1.0 M NaCl. Fractions with high purity were identified by SDS– PAGE and concentrated using a centrifugation filter with a 10 kDa cutoff (Amicon, Millipore). MBP-FUS-CHOP fusion type II and MBP-CHOP proteins were then flash frozen in 25% glycerol.

### Turbidity measurements

Turbidity was used to evaluate phase separation of 50 μM MBP-FUS-CHOP fusion type II and MBP-CHOP in the presence of 0.01 mg/mL TEV protease (~ 0.3 mg/mL in 50 mM Tris 1 mM EDTA 5 mM DTT pH 7.5 50% glycerol 0.1% Triton-X-100). The experiment was performed in 50 mM Tris 150 mM NaCl pH 7.4. Turbidity experiments were performed in a 96-well clear plate (Costar) with 70 μL samples sealed with optical adhesive film to prevent evaporation (MicroAmp, Thermo Fisher). The absorbance at 600 nm was monitored over time using a Cytation 5 Cell Imaging Multi-Mode Reader (BioTek) at 5 min time intervals for up to 12 h with mixing and subtracted from a blank buffer with no turbidity.

### DIC microscopy

For 50 μM MBP-FUS-CHOP type II fusion and MBP-CHOP, the samples were incubated with 0.03 mg/mL TEV protease for ~ 20 min before visualization. Samples were spotted onto a glass coverslip and droplet formation was evaluated by imaging with differential interference contrast on an Axiovert 200M microscopy (Zeiss).

## Acknowledgement

We thank Dennis McDaniel for his assistance with confocal microscopy. This project was supported by the National Institute of General Medical Sciences (Award Numbers R35GM119790 (to F.S.) and R01GM118530 (to N.L.F.)) and the National Institute of Neurological Diseases and Stroke (Award R01NS116176 (to N.L.F.)). The authors declare no competing financial interests.

## Author Contributions

Izzy Owen performed all in-cell experiments. Debra Yee designed and engineered transfections plasmids. Hala Wyne managed cell lines. Theodora Myrto Perdikari purified protein and performed in-vitro assays. Victoria Johnson purified protein. Izzy Owen, Frank Shewmaker and Nick L. Fawzi designed experiments and wrote the manuscript. Jeremy Smyth helped design imaging experiments. Robert Kortum helped with cancer cell lines.

## Abbreviations

(BRD4): Bromodomain containing protein 4
(C/EBP): CCAAT/Enhancer Binding Protein
(CHOP): CCAAT/Enhancer Binding Protein Homoglous Protein
(FRAP): Florescence recovery after photobleaching
(FUS): Fused in Sarcoma
(IDRs): Intrinsically disordered region
(LLPS): Liquid-liquid phase separation
(LC): Low complexity
(MBP): Maltose binding protein
(MLS): Myxoid Liposarcoma
(PrLD): Prion-like Domain
(STS): Soft tissue sarcoma
(SEs): Super Enhancers

